# Sequence-dependent model of genes with dual σ factor preference

**DOI:** 10.1101/2021.11.17.468920

**Authors:** Ines S. C. Baptista, Vinodh Kandavalli, Vatsala Chauhan, Mohammed N. M. Bahrudeen, Bilena L. B. Almeida, Cristina Palma, Suchintak Dash, Andre S. Ribeiro

## Abstract

*Escherichia coli* uses σ factors to quickly control large gene cohorts during stress conditions. While most of its genes respond to a single σ factor, approximately 5% of them have dual σ factor preference. The most common are those responsive to both σ^70^, which controls housekeeping genes, and σ^38^, which activates genes during stationary growth and stresses. Using RNA-seq and flow-cytometry measurements, we show that ‘σ^70+38^ genes’ are nearly as upregulated in stationary growth as ‘σ^38^ genes’. Moreover, we find a clear quantitative relationship between their promoter sequence and their response strength to changes in σ^38^ levels. We then propose and validate a sequence dependent model of σ^70+38^ genes, with dual sensitivity to σ^38^ and σ^70^, that is applicable in the exponential and stationary growth phases, as well in the transient period in between. We further propose a general model, applicable to other stresses and σ factor combinations. Given this, promoters controlling σ^70+38^ genes (and variants) could become important building blocks of synthetic circuits with predictable, sequence-dependent sensitivity to transitions between the exponential and stationary growth phases.

## 1. Introduction

In *E. coli*, genes are expressed by RNA polymerases (RNAP) core enzymes, which have 5 subunits (α_2_ββ’ω). When bound to a σ factor, they become able to recognize specific promoters and, from there, synthesize RNA [1]. Transcription is regulated mostly at the promoter regions, which typically harbor transcription factor (TF) binding sites (TFBS) and other regulatory sequence motifs [2-7]. This regulation is essential for cellular adaptability to both internal as well as external conditions [8,9].

Since σ factors are needed for transcription and because cells are able to regulate their numbers, they are themselves a regulatory mechanism of gene expression [4,10-13]. *E. coli* has seven different σ factors [14]. During exponential growth in optimal conditions, RNAP mostly transcribes genes with preference for σ^70^, responsible for basic cell functions [15]. Other σ factors are only present under specific stresses [16].

As an example, when the environment becomes depleted of components required for cell growth, *E. coli* usually switches to stationary growth, largely triggered by the appearance of RpoS (σ^38^), which activates ~10% of the genome [17,18], leading to key phenotypic modifications [18–24]. Meanwhile, since the concentration of RNAP core enzymes remains relatively constant [25] and since the RNAP numbers are lower than σ factor numbers, this will force σ factors to compete for RNAP’s [10,12,14,20,26,27]. Consequently, when σ^38^ numbers increase, the genes unresponsive to σ^38^, if previously active, will be indirectly negatively regulated [10,26,28].

For the σ factor regulatory system to be efficient, promoters need to have high specificity to one σ factor. In agreement, only a small fraction of promoters can recognize more than one σ factor [6,29,30]. Of these, the most common (84%) are the promoters responsive to both σ^70^ as well as σ^38^ [31], Supplementary Table S3 and Section 2.4.1). Following the workflow in Figure 1, we investigated how the promoter sequences recognized by both σ^70^ and σ^38^ (logos of positions -41 to -1 shown in Figure 2) relate to the dynamics of the genes that they control (named ‘σ^70+38^ genes’) prior, during, and following the transition to stationary growth.

**Figure 1.**
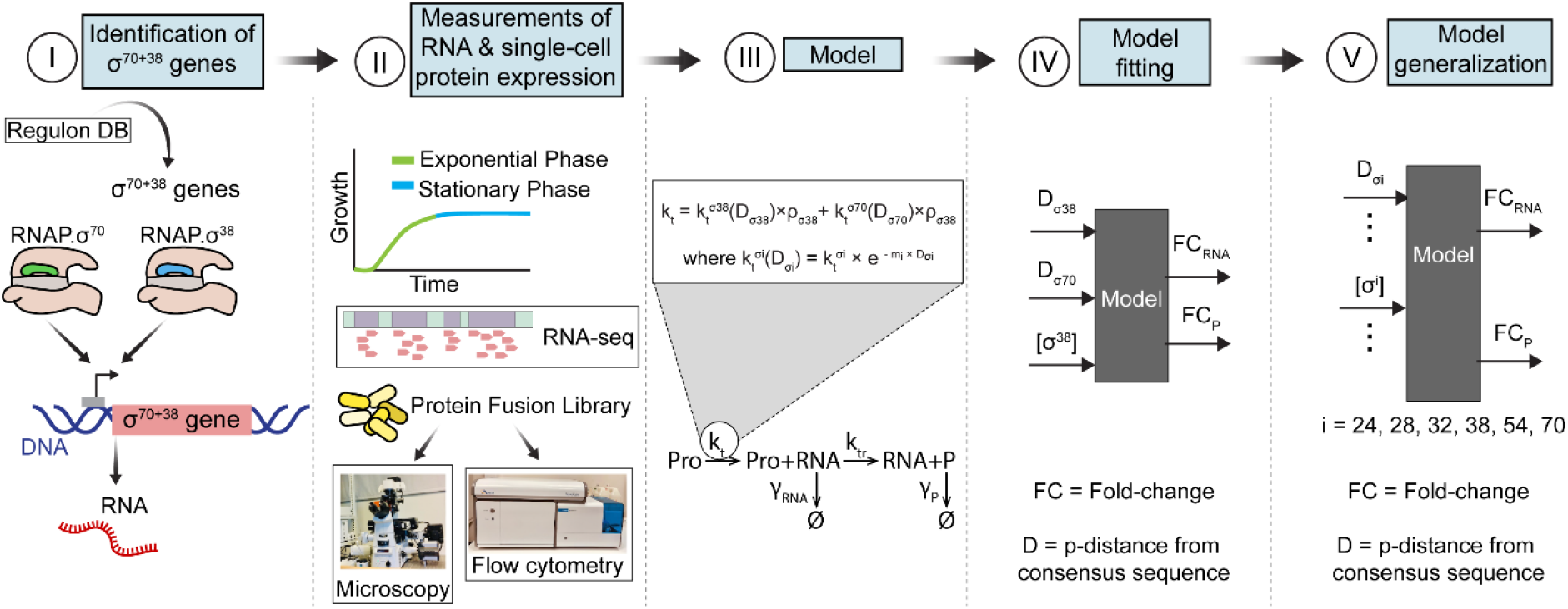
Workflow. **(I)** From Regulon DB [32], we identified genes controlled by single promoters with preference for both σ^70^ and σ^38^ (‘σ^70+38^ genes’). **(II)** Next, we measured RNA and single-cell protein levels of σ^70+38^ genes in the exponential and stationary growth phases. **(III)** Then, we proposed an empirically based model of gene expression fold changes of σ^70+38^ genes in RNA (FC_RNA_) and protein numbers (FC_P_) between growth phases. **(IV)** Afterwards, we tuned the model to fit how the promoter sequence affects the response to σ^38^. **(V)** Finally, we generalized the model to be applicable to genes with preference(s) for any set of σ factors.

**Figure 2.**
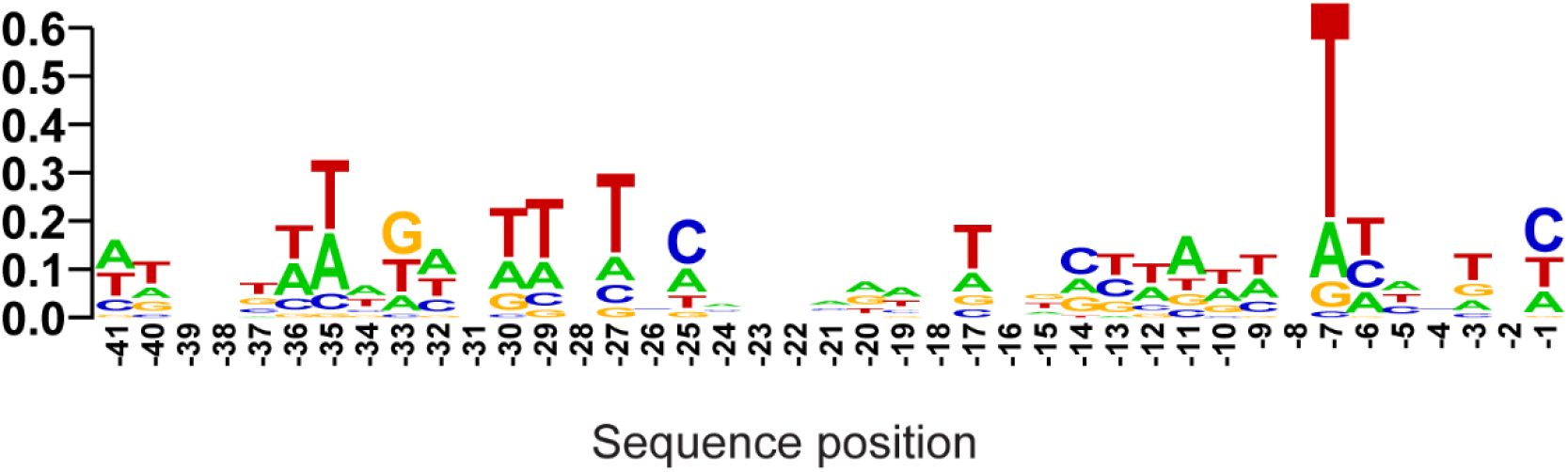
Sequence logo between positions -41 and -1 of promoter regions, with 0 being the TSS, of the 64 promoters with preference for both σ^70^ and σ^38^, obtained as described in Section 2.4.2. The logo of all other 8493 promoters in *E. coli* is shown in Supplementary Figure S1B.

## 2. Materials and Methods

### 2.1. Strains and media

*E. coli* strains and plasmids are listed in Supplementary Tables S1 and S2. We used YFP strains from the genetic stock center (CGSC) of Yale University, U.S.A [33] and, as support, a low-copy plasmid fusion library of fluorescent (GFP) reporter strains to track promoter activity [34]. In both, the fluorescent proteins under the control of the promoters of our genes of interest have been shown to be a good proxy for the native protein levels. For simplicity, we refer to these fluorescence levels as ‘protein levels’.

We used a RL1314 strain (rpoC::GFP) generously provided by Robert Landick (University of Wisconsin-Madison), to measure RNA Polymerase levels [35]. Their *rpoC* gene codes for β’ sub-unit endogenously tagged with GFP. Since *rpoC* codes for the β’ subunit, a limiting factor in the assembly of RNAP holoenzyme [36,37], its numbers serve as a good proxy for RNAP numbers. For simplicity, [RNAP] refers to the concentration of both RNAP core and holoenzymes in cells. Finally, we used a *MGmCherry* (*rpoS*::mCherry) strain to measure RpoS levels (kind gift from James Locke [19]). Their *rpoS* gene codes for σ^38^ endogenously tagged with mCherry. Finally, we used wild-type K12 MG1655 strain for control.

We used M9 medium (1xM9 Salts, 2 mM MgSO4, 0.1 mM CaCl2; 5xM9 Salts 34 g/L Na2HPO4, 15 g/L KH2PO4, 2.5 g/L NaCl, 5 g/L NH4Cl) supplemented with 0.2% Casamino acids and 0.4% glucose, and Luria-Bertani (LB) medium with 10 g peptone, 10 g NaCl, and 5 g yeast extract in 1000 ml distilled water. We used the antibiotics kanamycin and chloramphenicol from Sigma Aldrich, U.S.A.

### 2.2. Growth rate and growth phase

Growth rates were measured by spectrophotometry (BioTek Synergy HTX Multi-Mode Microplate Reader). From a glycerol stock (-80 °C), cells were streaked on LB agar plates (2%) and incubated at 37 °C, overnight. A single colony was picked, inoculated in LB medium with antibiotics (Section 2.1), and incubated at 30°C overnight with shaking. Overnight cultures were further diluted into fresh medium to an optical density of 600 nm (O.D._600_) of 0.01 and incubated for growth by shaking at 250 rpm at 37°C. OD_600_ was recorded every 20 min for 800 min. Cells were extracted at 150 min and 700 min after inoculation into fresh medium to represent cells in exponential and stationary growth phases, respectively (Figure 3A).

**Figure 3.**
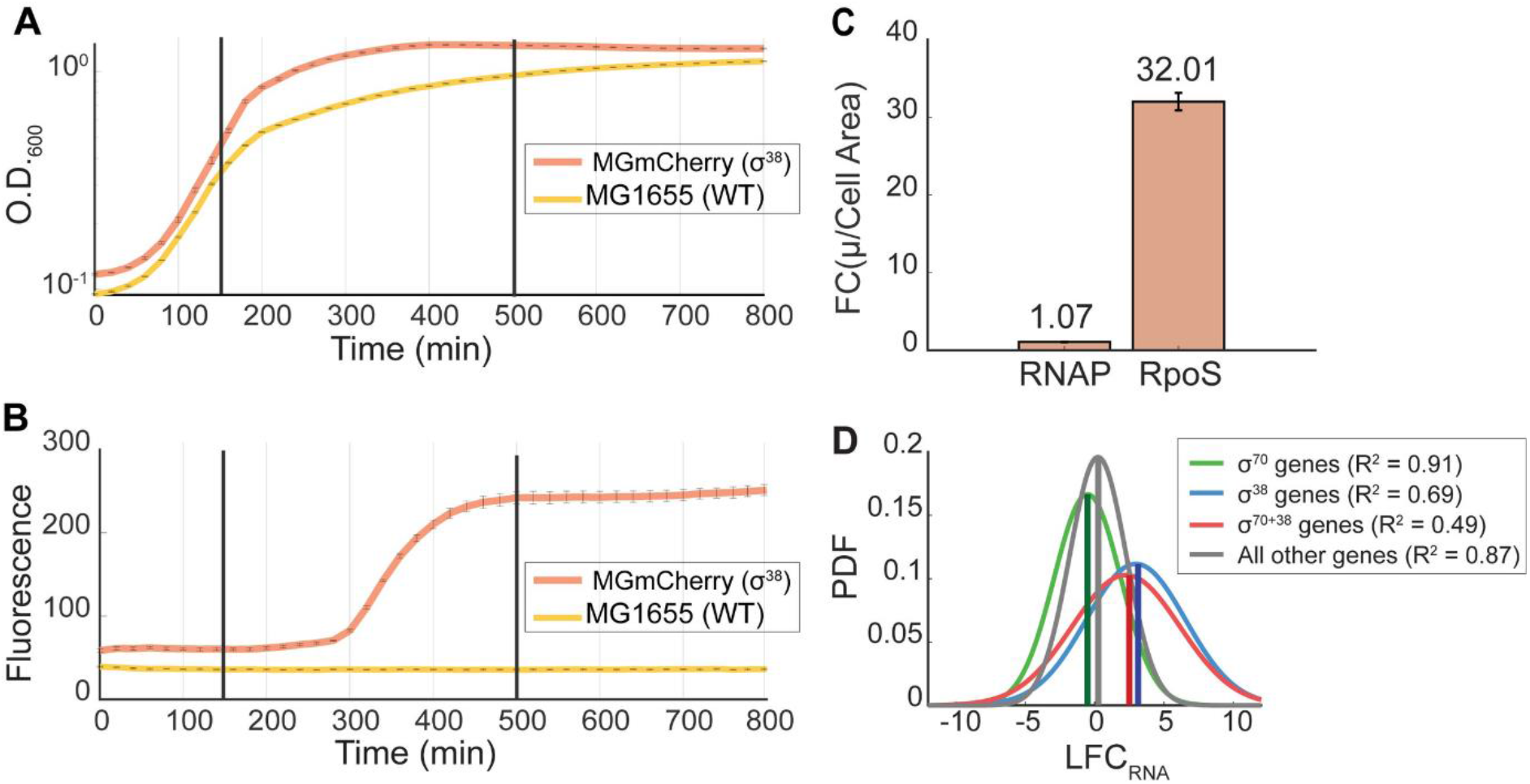
Cell growth, RNAP, RpoS and genome-wide RNA levels when changing from exponential to stationary growth. **(A)** Optical densities ‘O.D._600_’ of wild-type MG1655 (control) and MGmCherry (*rpoS::mcherry*) strains (Section 2.1). **(B)** Mean population fluorescence of WT and MGmCherry strains. Black vertical bars show the timing of measurements in exponential (150 min) and stationary (500 min onwards) growth. At 0 minutes, cells were moved to fresh medium (Section 2.2). **(C)** Fold-change between the exponential and stationary growth phases of mean cellular fluorescence relative to cell area (FC(μ/Cell Area)) due to changes in RNAP-GFP and in RpoS-mCherry, respectively (Cell Areas are shown in Supplementary Figure S3). Error bars are the standard error of the mean (SEM). **(D)** Gaussian fits to the distributions of LFC_RNA_ of gene cohorts. Vertical lines mark the mean. Given the high coefficient of determination (R^2^) of the fits, they likely capture well the shapes of the respective empirical distributions.

### 2.3. Gene expression measurements

To measure gene expression, we performed flow cytometry using an ACEA NovoCyte Flow Cytometer equipped with yellow (561 nm) and blue lasers (488 nm) and controlled by Novo Express V1.50 (Supplementary Section S1.1). Meanwhile, to measure σ^38^ *over time*, we used spectrophotometry (Supplementary Section S1.2). To convert protein expression levels to concentration levels, we used cell areas (as proxy for cell volume) which were obtained by microscopy and image analysis (Supplementary Section S1.3). Finally, we measured changes in RNA levels between growth conditions by RNA-seq (Supplementary Section S1.4).

### 2.4. Promoter sequences

#### 2.4.1. σ factor preference

Supplementary Table S3 informs on the genes’ σ factor preference. From Regulon DB v10.5 (August 14, 2019), we obtained lists of all transcription units (TUs), promoters (including σ factor preferences), and genes of *E. coli* [32]. Recently, we compared our lists with information obtained July 1, 2021 and found no changes that would affect the conclusions. TUs only differed by ~1%, promoters by ~0.5%, and genes by ~1%.

From the 3548 TUs (gene(s) transcribed from a single promoter), we extracted 2179 with known promoters, containing 2713 genes in total. To minimize interference to the classification of σ factor preferences arising from transcription by multiple promoters, we narrowed our list to 1824 genes transcribed by only one promoter and, of those, to the 1328 genes with known σ factor preference. From those, 1242 have a preference for only one σ factor, including 931 with a preference for σ^70^, and 93 with a preference for σ^38^. Conversely, 76 genes have promoters with a preference for two σ factors. Out of these, 64 are transcribed by only one promoter with a preference for σ^70^ and σ^38^.

#### 2.4.2. Sequence logo

Promoter sequence logos were created using WebLogo [38]. In each position (from -41 to -1), we counted how many times a nucleotide is present in all promoters considered. Then, we stacked all nucleotides (A, C, T, G) on top of each other, and sorted from the least found one in the bottom to the most present one on the top. For each position, we quantified its ‘bit’, as the difference between the maximum information possible (entropy given the 4 nucleotides) and the information considering the variability of the nucleotides (sum of the entropy for each of the 4 nucleotides) in that position (observed entropy): 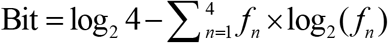. From this, the height of each letter is set to be proportional to its frequency of occurrence. Finally, the height of each position was normalized, so as to equal its corresponding amount of information (with the more conserved positions having more bits) [39].

#### 2.4.3. p-distance

The p-distance D of a promoter [40] is the fraction of its nucleotides between positions -41 to -1 (assuming that transcription start site starts at position +1) that differ from the consensus (most common) nucleotide in that position of a cohort of genes (here, genes with preference for a given σ factor). We extracted the consensus sequences related to each σ from RegulonDB [32]. To measure the D of promoters with preference for both σ^38^ and σ^70^, we calculated the p-distances D_σ38_ and D_σ70_, and then an overall p-distance, Δ, defined as: Δ = D_σ38_ – D_σ70_.

### 2.5. Statistical Analysis

We used two-sample Kolmogorov-Smirnov tests (KS tests) to compare distributions. Also, we used the Distribution Fitter app (MATLAB) to fit Gaussian fittings, the MATLAB’s curve fitting toolbox to fit curves, linear regression models to fit linear correlations, and least squares fitting [41] to fit Hill functions (Supplementary Section S1.6). To fit and validate surfaces, we used cross-validation resampling. Finally, we used Fisher’s exact tests to find overrepresentations in gene ontology (Supplementary Section S1.6).

## 3. Results

### 3.1. RNA fold changes when shifting to stationary growth

To study the dynamics of σ^70+38^ genes, we first identified when, after placing cells in fresh media, they transition from exponential and stationary growth (Section 2.2). We used both wild type (WT, control) cells and a MGmCherry strain [19] carrying fluorescently tagged σ^38^. From Figures 3A-C, cells are in the mid-exponential growth phase 150 min after moved to fresh medium, while their σ^38^ levels are low. Meanwhile, at 500 min onwards, cells are in stationary growth, while their σ^38^ levels are high. Meanwhile, the RNAP concentration was only ~7% higher during stationary growth (Supplementary Figures S3C-D).

We then performed RNA-seq, at 150 min and 500 minutes, and calculated the log_2_ fold changes in RNA levels between those moments (LFC_RNA_) (Section 2.3). In Figure 3D, we show Gaussian fits to the LFC_RNA_ distributions of σ^70+38^ genes, σ^70^ genes, and σ^38^ genes, as well as all other genes. In general, σ^70+38^ genes were nearly as upregulated as σ^38^ genes. On the other hand, σ^70^ genes were weakly downregulated, likely due to the expected indirect negative regulation [10,12,14,20,26,27]. Finally, most other genes (~2737 out of the 4308 genes) were relatively unresponsive.

### 3.2. Propagation of shifts in RNA numbers to protein numbers

Next, we measured single-cell protein levels (Section 2.3) of 9 out of the 64 σ^70+38^ genes (only 15 of the 64 are present in the YFP library (Section 2.1) and some of these exhibited too weak signals). These 9 genes, according to their LFC_RNA_, should cover most of the state space of response strengths of σ^70+38^ genes (Supplementary Section S1.5).

From measurements in exponential and stationary growth, we extracted means, μ_P_, and squared coefficient of variation, CV^2^_P_. We also calculated the log_2_ fold changes in μ_P_, LFC_P_. Since LFC_P_ and the corresponding LFC_RNA_ (Figure 3D) are linearly positively correlated, with a high R^2^ (Figure 4A), the changes in RNA levels of σ^70+38^ genes during the growth phase transition are propagating to their protein levels.

**Figure 4.**
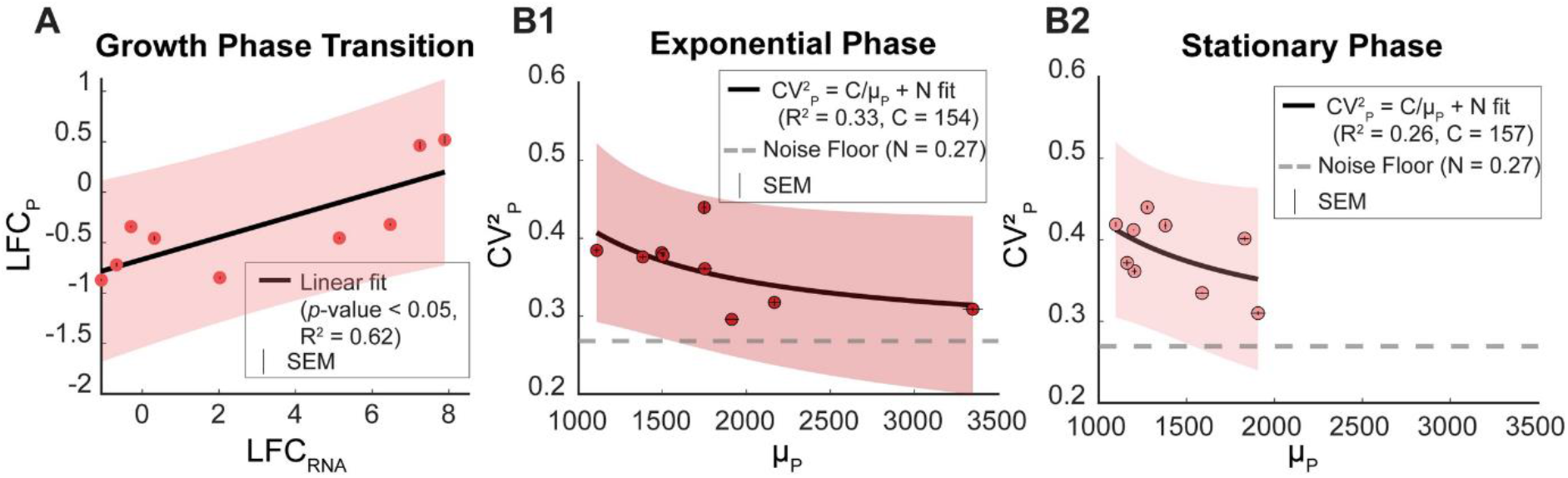
Single-cell protein levels of σ^70+38^ genes in the exponential and stationary growth phases. **(A)** Log_2_ fold-change of protein levels from exponential to stationary growth, LFC_P_, plotted against the corresponding LFC_RNA_. **(B1)** and **(B2)** CV^2^_P_ plotted against μ_P_ of σ^70+38^ genes in the exponential and stationary growth phases, respectively. Error bars (small) are the standard error of mean. All figures show the best fitting curves and their 95% confidence bounds (shadow areas). Horizontal dashed lines are the noise floors.

We then analyzed the variability in protein levels, prior and after the growth phase transition. In Figures 4B1 and 3B2, CV^2^_P_ decreases quickly with μ_P_ for small μ_P_, but slowly for high μ_P_. This is well described by a function of the form: 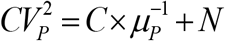 [33] (*N* is the noise floor, a lower bound on the cell-to-cell variability of protein levels in clonal populations due to extrinsic factors [42]), in agreement with [33,43,44].

### 3.3. Sequence-dependent model of σ^70+38^ genes

Based on the above, we proposed a sequence-dependent model for σ^70+38^ genes. We assumed that only σ^70^ is present in high numbers (in holoenzyme form) during the exponential phase and that only σ^38^ increases significantly in numbers when shifting to stationary growth, in agreement with [10,14,20,26,45–47].

The model is designed to account for competition between σ^70^ and σ^38^ to bind to RNAP core enzymes, since these exist in limited numbers [10,12,14,20,26,27]. For this, we set reactions for binding and unbinding of σ^70^ and σ^38^ to free floating RNAP core enzymes, (R1a) and (R1b), where *K^σ70^* and *K^σ38^* are the ratios between the respective association and dissociation rate constants:

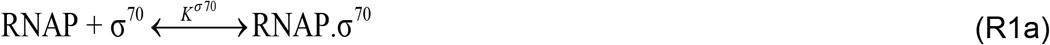

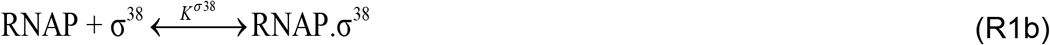

Given (R1a) and (R1b), the RNA production dynamics depends on the ratio between σ^70^ and σ^38^ numbers if, and only if, *K*^*σ*70^ and *K^σ38^* differ. The limited pool of RNAPs is accounted for since, from (R1a) and (R1b) alone:

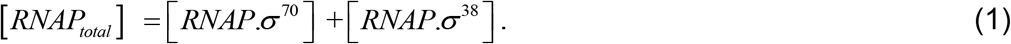

To introduce the dual responsiveness to σ^70^ and σ^38^, we defined two competing, sequence-dependent reactions of transcription, (R2a) and (R2b), differing in which holoenzyme binds to the promoter. The rates 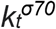 and 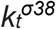 control the affinities to the holoenzymes and are expected to differ between promoters (and potentially with the transcription factors acting on the promoters, not represented for simplicity).

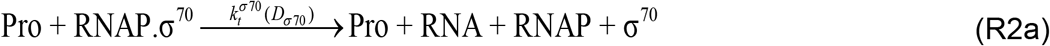

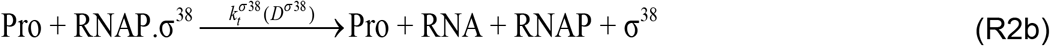

The reactions’ rate constants are set to be sequence dependent in what concerns σ^38^ and σ^70^ dependency. In detail, D_σ38_ and D_σ70_, are the p-distances, in nucleotides, of a promoter sequence from the consensus (average) sequence of promoters with σ^38^ and with σ^70^ dependency, respectively (Section 2.4.3):

Taken together, (R1a), (R1b), (R2a) and (R2b), model the transcription kinetics of σ^70+38^ genes before, during and, after shifting from exponential to stationary growth. The rates 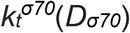 and 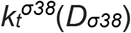 are dissected below.

### 3.4. Reduced model

The model above can be reduced since, first, the numbers of RNAP.σ^70^ and RNAP.σ^38^ in the cells are significantly larger than 1 [14,20,28,45-47]. As such, (R1a, R1b, R2a, and R2b) can be reduced to (R3a) and (R3b):

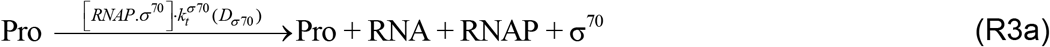

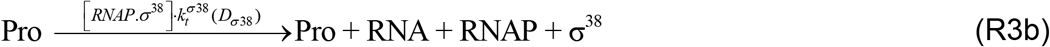

Further, one can merge (R3a) and (R3b) into a single transcription process (R4):

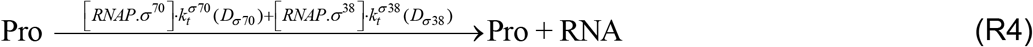

It was not possible to reduce the model further (e.g., by making *D_σ70_* a function of *D_σ38_*) since we failed to find evidence for a correlation between *D_σ70_* and *D_σ38_* (Supplementary Figure S6), except for the reduced set of σ^70+38^ genes *without* known input TFs (Figure 5C).

**Figure 5.**
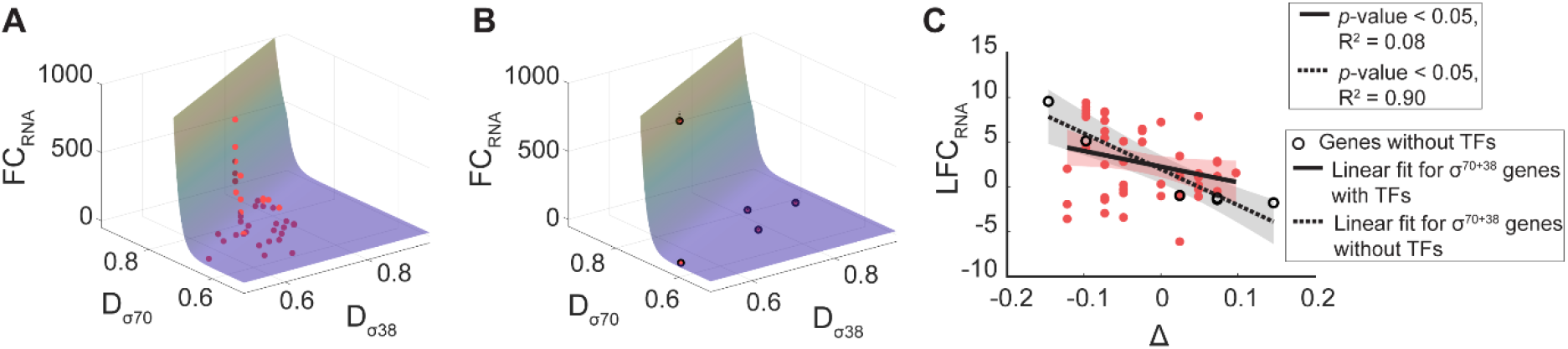
Model and measurements: fold changes in RNA levels of σ^70+38^ genes plotted against their promoter *p*-distances from the consensus sequence of σ^70^ dependent promoters (*D_σ70_*) and of σ^38^ dependent promoters (*D_σ38_*). **(A)** show the best fitting surfaces to FC(μ_RNA_) assuming an exponentially decreasing function. Only σ^70+38^ genes with input TFs and FDR < 0.05 are included. Light red points are above the surface, while dark ones are below. **(B)** Same plot but the surfaces are those obtained from (A1) and applied to σ^70+38^ genes without input TFs and FDR < 0.05. The dashed black lines depict the vertical distances between the estimated and measured FC(μ_RNA_). **(C)** Scatter plot of *Δ* = *D_σ38_* - *D_σ70_* plotted against LFC_RNA_. The shadows of the best fitting lines are the 95% confidence bounds.

Finally, we included reactions for translation of RNAs into proteins (R5), and for RNA (R6) and protein (R7) decay due to degradation and dilution by cell division:

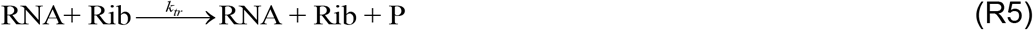

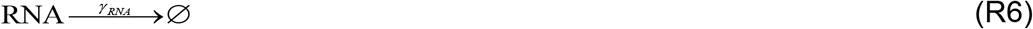

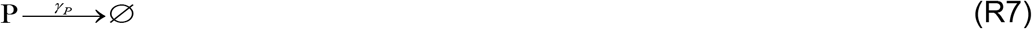

### 3.5. Analytical solution of the reduced model

Next, we obtained an analytical solution for the expected fold change in RNA numbers of a gene whose promoter has preference for both σ^70^ as well as σ^38^. From (R4), (R5), (R6), and (R7) the expected RNA numbers of a σ^70+38^ gene should equal:

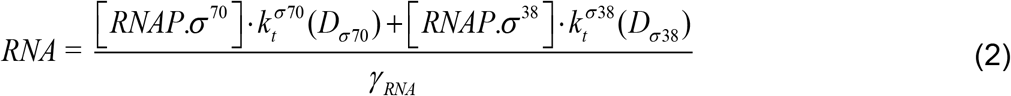

These rate constants are not expected to differ between the growth phases (as they depend on biophysical parameters, such as binding affinities, etc.). Consequently, the fold-change in RNA numbers between the growth phases should equal:

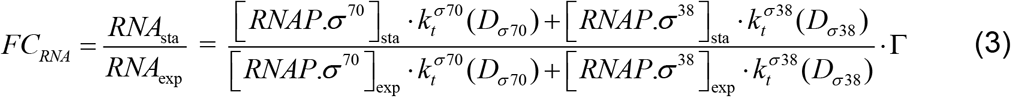

where Γ is the ratio between the RNA decay rates in the two growth phases:

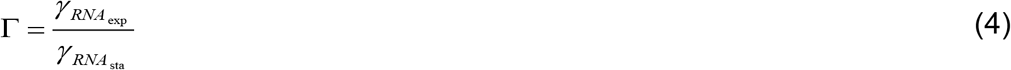

Since [*RNAP_total_*] is similar in the two growth phases (Figure 3C):

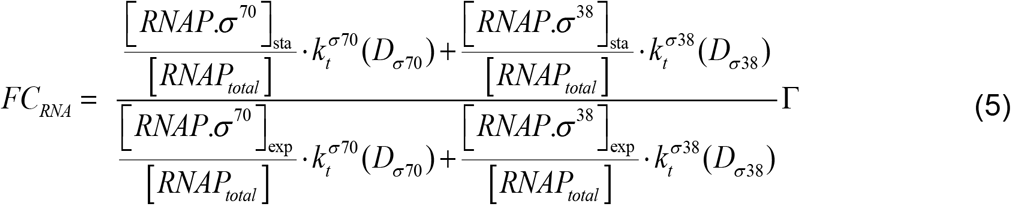

Next, considering equation (1), then:

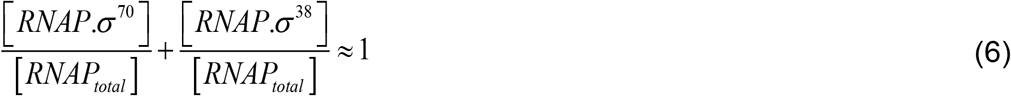

For simplicity, let *ρ_exp_* and *ρ_sta_* be:

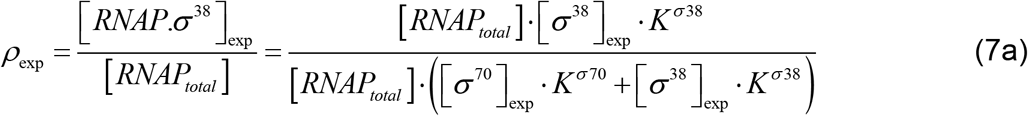

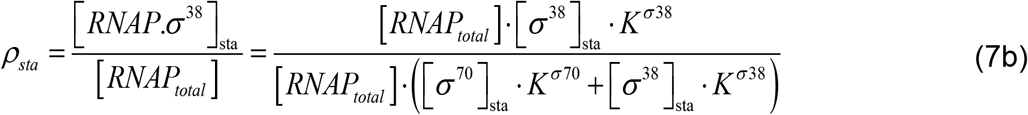

As such, from (5):

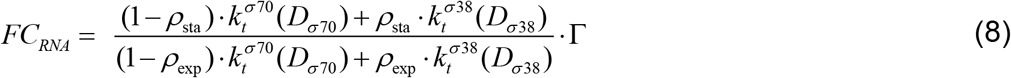

Finally, since in the exponential growth phase [*RNAP.σ^70^*]≫[*RNAP.σ*^38^], then *ρ*_exp_ ≈ 0. Thus, equation (8) can be simplified:

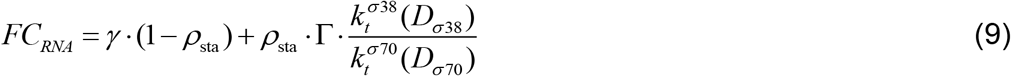

All parameters in equation (9) can be measured, except 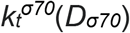 and 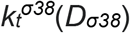. Meanwhile, in the stationary phase: [*σ*^38^]^sta^ = 0.3·[*σ*^70^] [20,26,46]. Also, *K^σ70^* = 5*K*^*σ*38^ [48]. Thus:

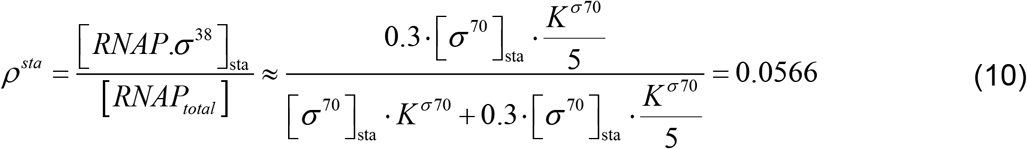

### 3.6. Promoter sequence affects the expression of σ^70+38^ genes

To model how promoter sequences (their *D_σ70_* and/or *D_σ38_*) influence the response strength of σ^70+38^ genes to σ^38^, we assumed basal transcription rates ‘*k_t0_^σ70^*’ and ‘*k_t0_^σ38^*’, when transcribed by RNAP.σ^70^ alone and by RNAP.σ^38^ alone, respectively.

To obtain the overall transcription rates of specific promoters, we then multiply *k_t0_^σ70^* and *k_t0_^σ38^* by a single-gene, sequence dependent function *f* to account for the influence of their *D_σ70_* and *D_σ38_*:

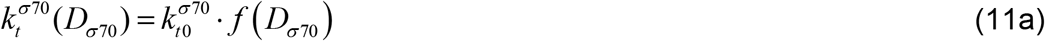

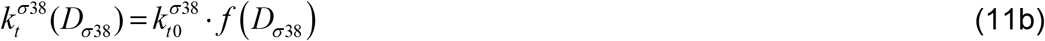

To define *f* we considered that, in general, as *D_σi_* increases, the promoter sequence should differ more from the ‘average’ sequence of promoters with preference for σ^i^. Thus, its affinity to σ^i^ should decrease. If this holds true, then the promoter consensus sequence of genes with preference for a given σ factor should have strong affinity to that σ factor. Thus, we hypothesized that the consensus sequence should have the strongest affinity. If true, it follows that as *D_σ38_* increases, the transcription rate by RNAP.σ^38^ decreases.

Having this, to model how the transcription rate of promoters recognized by σ^70^ and by σ^38^ specifically differs with *D_σi_*, we opted for an exponential function, *f*(*D_σi_*) = *e*^-m_i_.D_σi_^ where *i* = 38 or 70 and *m* is an empirical-based constant, similar to the one in [6,29,30] for σ^70^ genes. We first best fitted this function to σ^70+38^ genes *with* input TFs (Figure 5A), and then tested it if it could predict FC_RNA_ of σ^70+38^ genes *without* input TFs (Figure 5B). These genes should exhibit dynamics related to their promoter sequence (and thus the model), since are not subject to known TFs interference. From Table 1, we obtained an R^2^ of 0.98, suggesting that the exponential function can fit well the data (including when compared to other models (Supplementary Section S2.1)).

**Table 1.** Goodness of fit, measured by the R^2^ of the surface fitting of *FC_RNA_* as a function of *D_σ38_* and *D_σ70_*. Shown are the model and best fitting parameter values, where 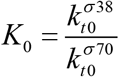. The surface was fitted to σ^70+38^ genes without input TFs and validated on genes σ^70+38^ genes with input TFs.

| Surface Equation | $FC_{RNA} = \gamma \cdot (1 - \rho_{sta}) + \gamma \cdot \rho_{sta} \cdot K_0 \cdot e^{-(m_{38} \cdot D_{\sigma 38} - m_{70} \cdot D_{\sigma 70})} \quad (12)$ | |
| --- | --- | --- |
| Coefficients | $m_{38}$ | 30.78 |
| | $m_{70}$ | 56.90 |
| | $\Gamma$ | 28.20 |
| | $K_0$ | $3.841 \times 10^{-9}$ |
| | $\rho_{sta}$ | 0.06 (Equation (10)) |
| $R^2$ | $\sigma^{70+38}$ genes without input TFs | 0.98 |

We further tested the model by randomly resampling (1000 times) the data from all σ^70+38^ genes into fitting and validation sets (Section 2.5). On average, the R^2^ to the testing sets equaled 0.5, which is only 30% lower than the R^2^ to the fitting sets. We find that the exponential model describes well the behavior of σ^70+38^ genes.

Next, from Figure 4A, the LFC of protein numbers (LFC_P_) correlates linearly with LFC_RNA_, as expected from the model (reactions R5 to R7) (Supplementary Section S2.2). From the best fitting line, we extracted a scaling factor, α (equaling 0.1), between LFC_RNA_ and LFC_P_, to be introduced in (13):

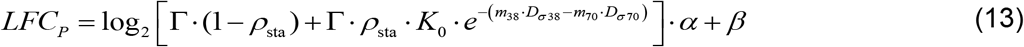

where β, equaling -0.67, is the intercept between the y axis and the best fitting line.

Finally, we observed that for σ^70+38^ genes without input TFs (previously used to validate the surface fitted to σ^70+38^ genes with TF inputs), *m_38_* and *m_70_* are similar (49.04 and 52.48, respectively), unlike for genes with input TFs (Table 1). Thus, for this restricted set of genes, we defined *Δ* = *D_σ38_* - *D_σ70_* (Section 2.4.3) and plotted again the respective LFC_RNA_ values. From Figure 5C, they correlate linearly.

We also searched for linear correlations between Δ and LFC_RNA_ in genes whose promoters have preference for σ^70^, σ^38^, σ^24^, σ^28^, σ^32^, or σ^54^ (Supplementary Figures S8A-F). No significant correlation with high R^2^ was found, including in the two small cohorts of genes whose promoters have preference for two σ factors, other than σ^70+38^ (Supplementary Figures S8G-H). Finally, we tried to fit the complete model (based on D_σ38_ and D_σ70_ separately) to these cohorts, but it was unable to fit well their behavior (Supplementary Table S6).

### 3.7. Expanding the model to the growth phase transition period

The model above was designed to be applied in either growth phase. Here, we expanded it to be applicable to the transition period between the growth phases, assuming σ^38^ to be the main regulatory molecule. We collected temporal data on protein numbers of three σ^70+38^ genes, specifically pstS, aidB and asr, selected to represent σ^70+38^ genes with strong, mild, and weak response strengths to the growth phase transition, respectively (Figure 6A).

**Figure 6.**
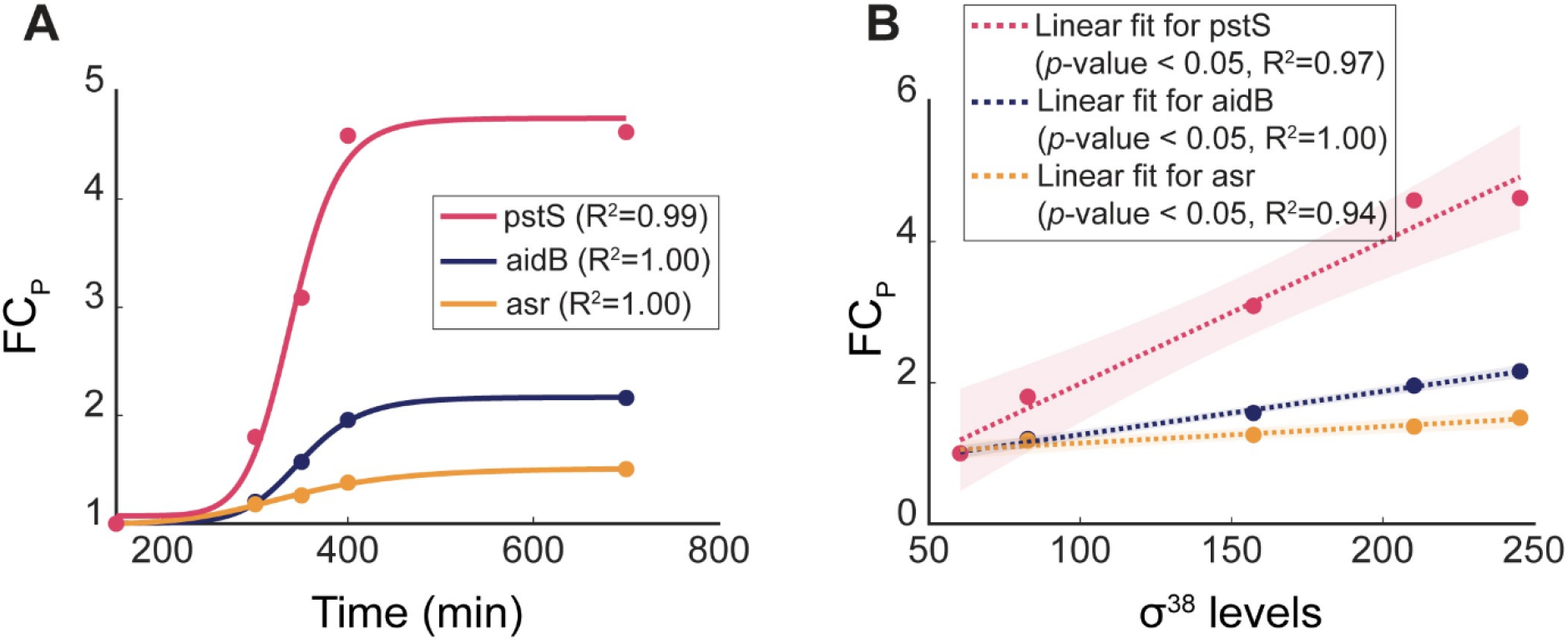
Temporal changes in the fold-change of protein levels of σ^70+38^ genes as σ^38^ changes. **(a)** Protein levels of three σ^70+38^ genes prior to, during, and after the transition from the exponential to the stationary growth phase. The balls are the empirical mean fold changes (FC_P_) in protein expression levels relative to the first-time moment. The lines are the corresponding best fitting Hill functions (parameter values in Supplementary Table S5). **(b)** Scatter plot of FC_P_ against the corresponding σ^38^ levels (data from Figure 3B) over time. The shadows are the 95% confidence bounds.

Their empirical data was plotted in Figure 6A. Then, we applied Hill functions (given the data on σ^38^ from Figure 3B), which fitted well the data. Moreover, we found a linear relationship between the fold changes in protein levels of the 3 genes, and the σ^38^ levels over time (Figure 6B).

The dependency on σ^38^ levels in model is set by *ρ_sta_* which is a function of the time-dependent σ^38^ levels (equation (7b)). Thus, given the goodness of fit of the Hill functions, we set the following time-dependent model:

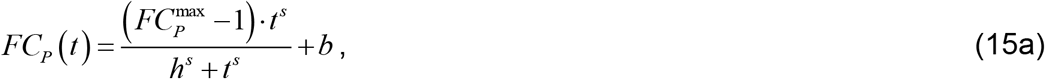

where:

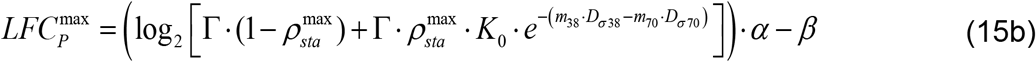

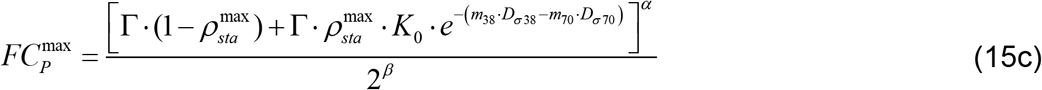

Here, 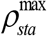 is the final (thus, maximum) concentration of RNAP.σ^38^ relative to the total concentration of bound RNAPs. Meanwhile, 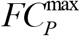 is the expected fold change in protein numbers of a σ^70+38^ gene, which is reached after completed the transition to stationary growth. Finally, *b* (the intercept) is the expression level controlled by the promoter of interest, when in the exponential growth phase. Meanwhile, *s* (the slope) is the response strength, and *h* (the half-activation coefficient) is a measure of the response timing to changes in σ^38^ levels.

### 3.8. Model generalization

Since we found a correlation between the promoter sequences of σ^70+38^ genes and their response to σ^38^, it may be that genes whose promoters have different σ preferences will exhibit, in some cases, similar sequence-dependent behaviors during the stresses that they respond to. Thus, we generalized the model to be applicable to any stress and responsive gene cohort.

As a general example, we set a model for genes responsive to all seven σ factors of *E. coli*, by expanding reactions (R1a) and (R1b) to 7 reactions as follows:

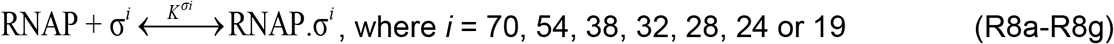

Given this, we generalize eq. (1) to account for all the σ factors in holoenzyme form:

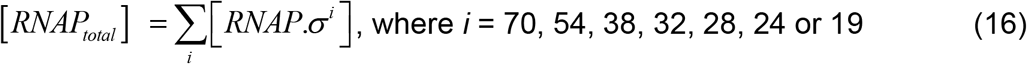

Finally, we generalize (R4) as follows:

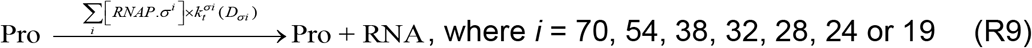

This general model can be tuned based on the numbers of each σ factor present in the conditions considered, and the consensus sequences to each σ factor (Supporting Table S2). In addition, following the findings in Section 3.7, it should be feasible to introduce factors to account for the timing of the changes in the transition period.

## 4. Discussion

Past studies have identified differences between σ^70+38^ and σ^38^ genes (e.g., Regulon DB sets this classification from ChIP-Seq and other data [32]). We additionally found a small difference in response kinetics to the growth phase shift (Figure 2D). Also, their sequence logos differ, particularly their consensus level in the ~ -35 region (Figure 1 and Supplementary Figure S1A). Finally, their ontologies differ with σ^70+38^ genes being more involved in respiration (Supplementary Table S4A), while σ^38^ genes are more involved in metabolic processes (Supplementary Table S4B).

From their sequence and dynamics, we proposed and validated a promoter sequence-dependent kinetic model of genes controlled by promoters responsive to both σ^70^ and σ^38^. The model, which accurately predicts how the dynamics changes from exponential and stationary growth, is an expansion of a past model of promoters with preference for σ^70^ alone [26]. However, it has two competing reactions for transcription by RNAP when bound to σ^70^ and when bound to σ^38^, respectively. In addition, these two reactions’ rate constants are sequence-dependent, in line with the hypothesis that the consensus sequence of promoters with preference for a σ factor should provide the strongest affinity to that σ factor (in general). Further, the model is applicable during growth phase transition, based on the σ^38^ level at any given moment.

The identification of a correlation between the p-distances of the promoter sequence of σ^70+38^ genes and their response strength to the growth phase transition is, to our knowledge, a unique feature. So far, sequence dependent behaviors have not been reported for any cohort of natural promoters of *E. coli*, even under stable growth conditions. Similar relationships have only been reported for synthetic libraries of variants of single promoters (thus, sequence-restricted) [5,6,30,49].

Since it remains challenging to predict if a sequence can act as a promoter and, if so, with which strength and under what regulatory mechanisms [5], the observed predictability of the dynamics from the sequence for the natural cohort of σ^70+38^ genes is of interest for three main reasons. First, these promoters and their variants (with varying p-distances from the consensus sequences of σ^70^ and σ^38^ genes) could become key components of future circuits whose predictable kinetics. Further, the circuits would likely be functional in both the exponential and stationary growth phase, with tunable responsiveness to the phase transition (given the results in Figure 6). Finally, this approach could become a starting point for broader models of natural promoters with a sequence-predictable adaptability to stresses.

However, it may prove difficult to find similar relationships between the sequences of other natural promoters and their responsiveness to other σ factors (or global regulators). E.g., we failed to find a relationship between the sequences of promoters responsive to σ^38^ alone and their response strength to σ^38^ (even for genes without TF inputs (Supplementary Figure S8B)).

It may be possible to expand this model in various ways. First, it may be applicable to genes responsive to σ^70^, σ^54^, σ^32^, σ^28^, σ^24^, or dual combinations, when the respective σ factors change numbers. Also, it should be possible to consider the influence (interference or enhancement) of TFs in the genes’ responsiveness to the σ factor.

While it is expected that the model should be applicable to various bacteria, it may be further applicable to eukaryotes. Namely, it is plausible that similar mechanisms of dual global regulation exist in some eukaryotic promoters that depend on TFIID complexes, which are highly conserved [50–52]). The transcription of these promoters only initiates after a TFIID factor binds to their TATA box. Such TFIID factors, in addition to a TBP (TATA binding protein), also require one TAF (TBP associated factor). Since eukaryotic cells have several different TAFs, this mechanism has similarities to σ factor regulation [53]. In the future, it should be interesting to explore the applicability of our model to such eukaryotic promoters responsive to multiple TAF [54].

From the biological point of view, first, it should be interesting to investigate if the sequence-dependent responsiveness of σ^70+38^ genes to σ^38^ levels could explain why their promoters (from positions -41 to -1) are highly conserved (Figure 2). Another potential reason for why they are conserved could also that the TFs that they code for commonly serve as input TFs to essential genes [55] (2.5 times more than by chance, Fisher test *p*-value < 0.05).

Finally, over-representation tests of the ontology [56,57] of these genes suggest that they are commonly involved in respiration (Supplementary Table S4A). In agreement, respiration is highly affected when changing from exponential to stationary growth [23,58], since aerobic respiration is reduced, while fermentation and anaerobic respiration are enhanced [18].

Moreover, of the genes associated with these biological functions, σ^70+38^ genes are amongst the most responsive to the growth phase transition (Supplementary Figure S4). This suggests that they may control the most altered processes during the adaptation. Therefore, externally regulating σ^70+38^ genes may give significant control over these processes. This is particularly appealing since the control could be exerted by promoter sequence modifications (i.e., tuning p-distances) with largely predictable effects. This strategy could contribute to the engineering of synthetic circuits with tailored responses to growth phase transitions.

## Supplementary data

### Supplementary Material

Supplementary Material.

**Supporting Table S1.** List of genes classified as σ^70^, and σ^38^ or as both σ^70^ and σ^38^ dependent.

**Supporting Table S2.** Consensus sequences of promoters with preference for one σ factor.

**Supporting Table S3.** Fold changes in RNA levels of genes with a promoter with preference for σ^70^ and σ^38^. Measurements by RNA-seq in the exponential and stationary growth phases.

**Supporting Table S4.** Statistics of single-cell distributions of fluorescence of cells measured by flow cytometry.

## Acknowledgements

The authors acknowledge the Tampere facility of Flow Cytometry for their service.

