## Supplementary Material for "Sequence-dependent model of genes with dual σ factor preference"

### S1. Supplementary Materials and Methods

#### S1.1. Flow cytometry

Cells were diluted 1:10000 into 1 ml of PBS vortexed for 10 sec. For each gene and condition, we performed 3 biological replicates, acquiring 50,000 events for each. For GFP and YFP, we used the blue laser for excitation and the FITC-H channel (520/20 nm filter) for emission. For mCherry, we used a yellow laser for excitation and PE-Texas Red (615/20 nm filter) for emission. We collected events at a flow rate of 14 µl/minute, a core diameter of 7.7 µM, and adjusted PMT voltage for each parameter. We have set the FSC-H detection threshold to 5000.

We collected single-cell fluorescence distributions. To remove outliers due to debris, cell doublets, and other undesired events, we performed gating by setting the maximum values of SSC-H to $5\times10^4$ and of FSC-H to $10^5$. These limits allowed observing the whole distributions, without cutting their natural tails. Finally, we discarded “far out” events using Tukey’s fences [1], which had little effect on the results.

Similarly, the background fluorescence ended up not being subtracted, since it did not affect the results (Supplementary Figure S2). This was tested by subtracting the background fluorescence from flow cytometry data as in [2,3], by applying the equation (S1) [3]:

$$M_p = M_T - M_{cell} \qquad \text{(S1)}$$

where $M_p$ is the real mean cell fluorescence from the fluorescent protein of interest, $M_T$ is the measured mean cell fluorescence, and $M_{cell}$ is the mean cell auto-fluorescence. Similarly, to correct the variance ($\sigma^2$) of the measured fluorescence, one could apply equation (S2) [3]:

$$\sigma_p^2 = \sigma_T^2 - \sigma_{cell}^2 \qquad \text{(S2)}$$

From (S1) and (S2), we derived equation (S3), to correct the squared coefficient of variation, $CV^2$, of the single-cell protein expression levels.

$$CV_p^2 = \left( \frac{\sigma_p}{M_p} \right)^2 \qquad \text{(S3)}$$

### S1.2 Spectrophotometry

Time-lapse protein fluorescence of MG*mCherry* cells was measured by a BioTek Synergy HTX Multi-Mode Microplate Reader. Overnight cultured cells were diluted 1:1000 times into fresh M9 medium, aliquoted into 24 well dark bottom microplates, and kept at 37°C with shaking of 250 rpm. Fluorescence intensities were recorded for 14 hours, every 20 min, using excitation (590/20 nm) and emission (645/20 nm) filters. Temporal profiles of fluorescence intensity were extracted using Gen5 software, based on 6 biological repeats.

### S1.3 Microscopy and Image analysis

To collect microscopy data, cells were placed between a coverslip and agarose gel pad (2%) and visualized by a confocal laser-scanning system, using a 100× objective. Green fluorescence images were captured using a 488 nm laser and a 514/30 nm emission filter. Phase-contrast images were simultaneously acquired for segmentation and to assess health, morphology, and physiology.

Images were analysed using the “CellAging” software [4]. After automatically segmenting cells and manually correcting errors, we applied 2D Gaussian filters to remove measurement noise and extracted each cell’s total fluorescent intensity.

### S1.4 RNA-seq

#### *Sample preparation*

Cells were grown until reaching either the exponential or the stationary growth phase. At each of these moments, they were treated with a double volume of RNA protect bacteria reagent (Qiagen, Germany) for 5 min at room temperature, to prevent RNA degradation. Cells were then pelleted and frozen immediately at -80 ºC. After unfreezing the cells, total RNA was extracted using RNeasy mini-kit (Qiagen) according to the instructions. RNA was treated twice with DNase (Turbo DNA-free kit, Ambion, Life Technologies, U.S.A.) and quantified using Qubit 2.0 Fluorometer RNA assay (Invitrogen, Carlsbad, CA, USA). Quality of total RNA was determined by gel electrophoresis, using 1% agarose gel stained with SYBR safe (Invitrogen, U.S.A.). RNA was detected using UV with a Chemidoc XRS imager (BioRad, U.S.A.).

#### *Sequencing*

Sequencing was performed by GENEWIZ, Inc. (Leipzig, Germany). The RNA integrity number (RIN) of the samples was obtained with Agilent 4200 TapeStation (Agilent Technologies, Palo Alto, CA, USA). Ribosomal RNA depletion was performed using Ribo-Zero Gold Kit (Bacterial probe) (Illumina, San Diego, CA, USA). RNA-seq libraries were constructed using NEBNext Ultra RNA Library Prep Kit (NEB, Ipswich, MA, USA). Sequencing libraries were multiplexed and clustered on 1 lane of a flow-cell. Samples were sequenced using single-index, 2x150bp paired-end (PE) configuration on an Illumina HiSeq instrument. Image analysis and base calling were conducted with HiSeq Control Software (HCS). Raw sequence data (.bcl files) were converted into fastq files and de-multiplexed using Illumina bcl2fastq v.2.20. One mismatch was allowed for index sequence identification.

***Data analysis***

RNA-seq data analysis pipeline: i) RNA sequencing reads were trimmed to remove adapter sequences and nucleotides with poor quality with Trimmomatic [5] v.0.39. ii) Trimmed reads were then mapped to the reference genome (*E. coli* MG1655, NC_000913.3) using the STAR v.2.5.2b aligner [6]. iii) *featureCounts* from Rsubread R package (v.1.34.7) was used to calculate unique gene hit counts [7]. Genes with less than 5 counts in more than 3 samples, and genes whose mean counts are less than 10, were removed. iv) Next, the unique gene hit counts were used for subsequent differential expression analysis, using DESeq2 R package (v.1.24.0) [8] to compare gene expression between samples and calculate p-values and log2 of fold changes (LFC) using Wald tests (function *nbinomWaldTest*). P-values were adjusted for multiple hypotheses testing (Benjamini–Hochberg, BH procedure [9]) and genes with adjusted p-values (False discovery rate (FDR)) smaller than 0.05 were classified as differentially expressed (D.E.).

Due to filtering (iii), only 783 (777 D.E.) out of the 931 $\sigma^{70}$ genes, 87 (all D.E.) out of the 93 $\sigma^{38}$ genes, 57 (56 D.E.) out of the 64 $\sigma^{70+38}$ genes, and 3257 (2737 D.E.) out of 3607 remaining genes were assessed by RNA-seq. Other genes did not produce enough reads.

### S1.5. Genes chosen to measure single-cell protein abundance

The genes selected for flow cytometry have a fold-change in protein levels which correlates significantly with the fold-changes in RNA levels (Figure 4A, main manuscript). However, to be good representatives, these genes should largely cover widely and homogeneously the state space of the RNA dynamics of the cohort of all genes. To test this, we performed a KS test (Section 2.5, main manuscript) to compare the $LFC_{RNA}$ distribution of the selected genes with the distribution of the entire cohort. The test did not reject that the two distributions are from the same distribution (*p*-values > 0.05). As such, we conclude that the sub-cohort is a good representative of the full cohort.

Next, to verify if the sub-cohort represents well the state space of protein numbers in *E. coli*, we compared $\mu$ and $CV^2$ of each of the genes selected against the remaining genes (Supplementary Figure S5A, data from [10]). KS tests for $\mu$ and $CV^2$ showed that neither null hypothesis is rejected (*p*-values > 0.05).

Finally, the set of genes selected also largely covers widely and homogeneously the state space of the promoter sequences of all $\sigma^{70+38}$ genes with the same preferences for $\sigma$ factors (Supplementary Figures S5B-C). The null hypothesis of the KS test between this sub-cohort and the cohort of all $\sigma^{70+38}$ genes was not rejected (*p*-values > 0.05) for both $D_{\sigma38}$ and $D_{\sigma70}$.

#### S1.6. Statistics

In detail, data points from microscopy, flow cytometry, etc. are from at least 3 biological replicates each. Meanwhile, to assess if genes measured by flow cytometry were a good representative of the set of all $\sigma^{70+38}$ genes, we used two-sample Kolmogorov-Smirnov tests (KS tests) to compare their sequences ($D_{\sigma38}$ and $D_{\sigma70}$) and dynamics ($LFC_{RNA}$, μ, and $CV^2$).

Gaussian fittings were made by the Distribution Fitter app (MATLAB). To evaluate its fitting, we calculated the coefficient of determination ($R^2$) between the PDF of the distributions and the PDF of the fitting. Linear correlations were estimated by least-squares fitting (FITLM of MATLAB) and considered significant if the *p*-value of an *F*-test on the fitted regression line was smaller than 0.05. Finally, other curves were fitted using MATLAB's curve fitting toolbox (apart from the fitting of Hill functions, which was done using the Hill function [11]).

To fit and validate surfaces relating the promoter sequence (p-distance) to the RNA fold-change, we used a cross-validation resampling method to unbiasedly define training and test sets for creating and validating surfaces, respectively ('cvpartition' in MATLAB). In each of the 1000 iterations, we randomly partitioned the data points into 2 complementary subsets ($d_1$ and $d_2$). Next, the surfaces were fitted to the $d_1$ set, and tested against $d_2$ by calculating the $R^2$. After, the opposite is done. Finally, we filtered the $R^2$ values for all fitting and validating sets, to remove those whose $R^2$ values for both the fitting and validating sets were not between 0 and 1.

For gene ontology representations, we performed overrepresentation tests using PANTHER Classification [12] to find significant overrepresentations by Fisher's exact tests. For *p*-values < 0.05, the null hypothesis that there are no associations between the gene cohort and the corresponding GO of the biological process were rejected. This *p*-value was corrected by calculating if the overall False Discovery Rate (FDR) is < 0.05. FDR was calculated by the Benjamini-Hochberg procedure [9].

### S2. Supplementary Results

#### S2.1. Comparison of models relating sequence and RNA dynamics

To model how a promoter sequence controls the transcription kinetics of promoters recognized by $\sigma^{70}$ and by $\sigma^{38}$, in addition the exponential model, we also considered a linear model, $f(D_{\sigma i}) = 1 - m_i \cdot D_{\sigma i}$, and a rational model, $f(D_{\sigma i}) = \frac{1}{1 + m_i \cdot D_{\sigma i}}$ (transcription rate inversely proportional to $D_{\sigma i}$), where $i$ = 38 or 70 and $m$ is an empirical-based constant.

We fitted the models (Supplementary Figure S7) to the empirical fold changes (FC) of $\sigma^{70+38}$ genes ($FC_{RNA}$) with known input TFs (Supporting Table S3) (FC rather than LFC were used, because this allowed for simpler equations for the fitting surfaces). We found that these models did not fit the data better than the exponential model (i.e. their $R^2$ values were lower).

#### S2.2. Expected log fold changes in protein numbers

From reactions R5 to R7 in the main manuscript, the steady-state solution of the number of proteins P is:

$$P = \frac{k_{tr} \cdot Rib \cdot RNA}{\gamma_P} \qquad \text{(S4)}$$

Therefore, the fold-change in protein expression, $FC_P$, when cells transition from the exponential to the stationary growth phase is:

$$FC_P = \frac{P_{\text{sta}}}{P_{\text{exp}}} = \frac{\left(\dfrac{k_{tr}^{\text{sta}} \cdot Rib_{\text{sta}} \cdot RNA_{\text{sta}}}{\gamma_P^{\text{sta}}}\right)}{\left(\dfrac{k_{tr}^{\text{exp}} \cdot Rib_{\text{exp}} \cdot RNA_{\text{exp}}}{\gamma_P^{\text{exp}}}\right)} = \frac{k_{tr}^{\text{sta}} \cdot Rib_{\text{sta}} \cdot \gamma_P^{\text{exp}}}{k_{tr}^{\text{exp}} \cdot Rib_{\text{exp}} \cdot \gamma_P^{\text{sta}}} \frac{RNA_{\text{sta}}}{RNA_{\text{exp}}} = \Omega \cdot FC_{RNA} \qquad \text{(S5)}$$

where $\Omega = \dfrac{k_{tr}^{\text{sta}} \cdot Rib_{\text{sta}} \cdot \gamma_P^{\text{exp}}}{k_{tr}^{\text{exp}} \cdot Rib_{\text{exp}} \cdot \gamma_P^{\text{sta}}}$, and $FC_{RNA}$ is given by equation 9 in the main manuscript.

Converting equation (S5) to a logarithmic scale to match the $LFC_{RNA}$ data from RNA-seq, we get:

$$LFC_P = \log_2\left(\Omega\right) + LFC_{RNA} \qquad \text{(S6)}$$

Given this, a linear fit in a plot of the LFC in protein numbers against the LFC in RNA numbers should have a slope of 1. However, this is not the case (Figure 4A in the main manuscript).

This has (at least) two causes. First, in real cells, if the RNA numbers are very small or very large, this linearity is not expected because of errors in translation, limited numbers of ribosomes [13], etc. Second, the RNA-seq and flow cytometry data are not expected to have a 1 to 1 relationship since these techniques differ in sensitivity, etc.

Consequently, the relationship (S6) needs to be corrected by multiplying a scaling factor, named ‘α’.

### S3. Supplementary Figures

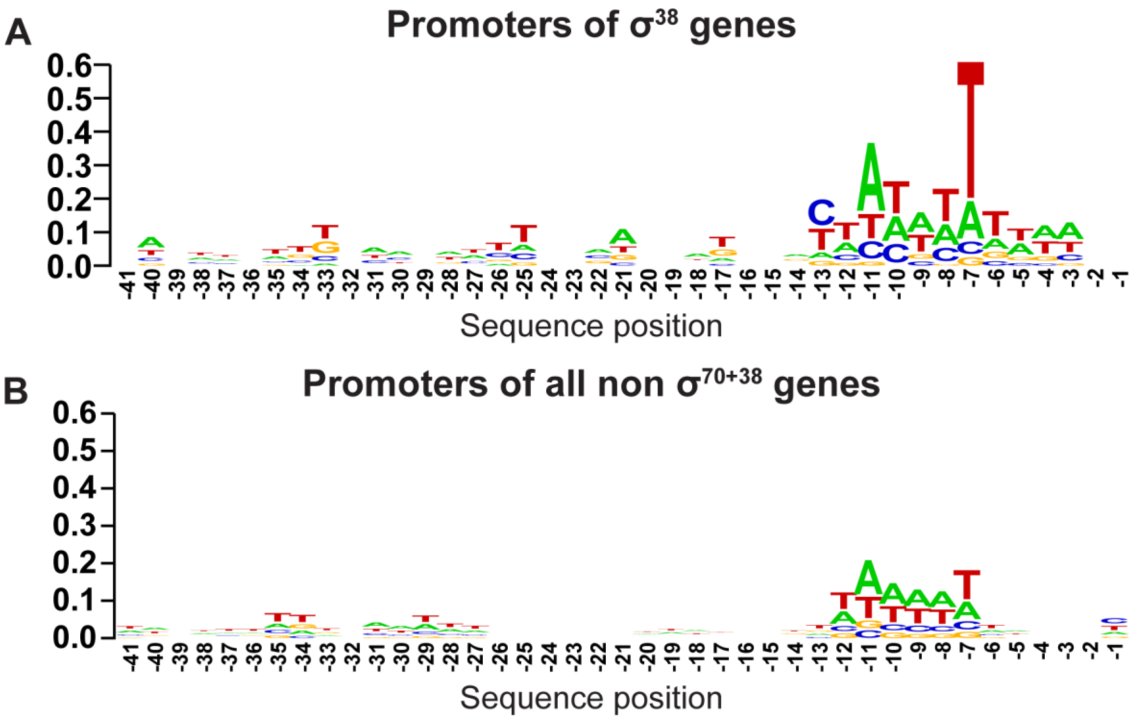


**Figure S1**. **(A)** Sequence logos between positions -41 and -1 of promoter regions, with position 0 being the TSS, of the 93 promoters with preference for $\sigma^{38}$; and **(B)** of the 8493 promoters in *E. coli* that do not have preference for both $\sigma^{70}$ and $\sigma^{38}$.

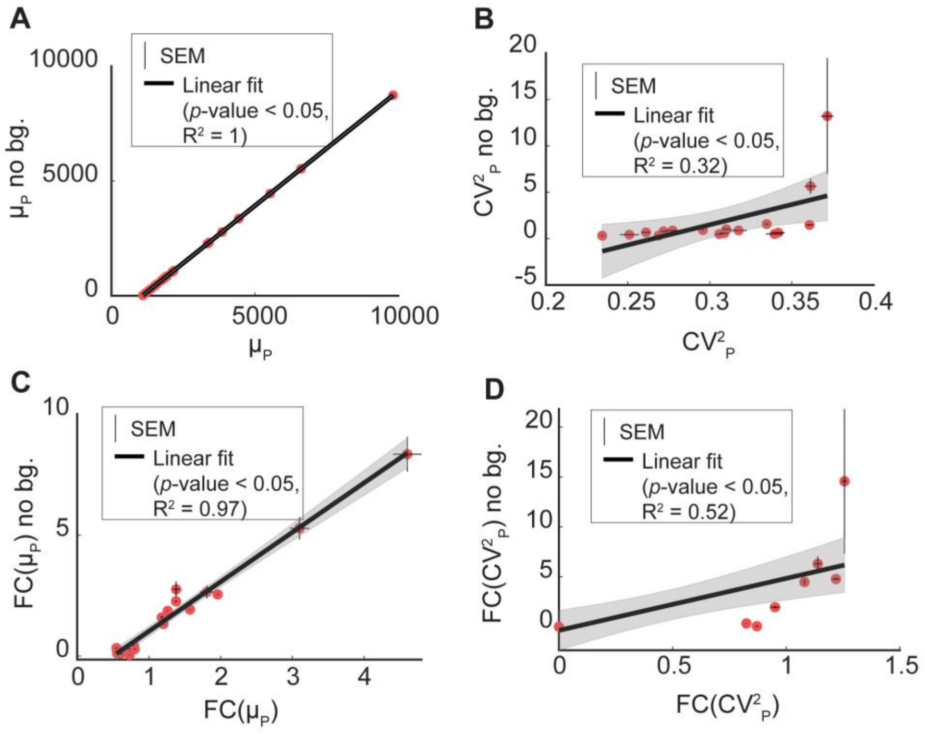


**Figure S2**. Effects of removing the background fluorescence (bg) on the **(A)** mean single-cell fluorescence ($\mu_P$); **(B)** $CV^2$ of the single-cell fluorescence ($CV^2_P$); **(C)** fold-change of the single-cell fluorescence ($FC(\mu_P)$) and **(D)** fold-change of $CV^2$ of single-cell fluorescence ($FC(CV^2_P)$). The shadows of the best fitting lines are the 95% confidence bounds.

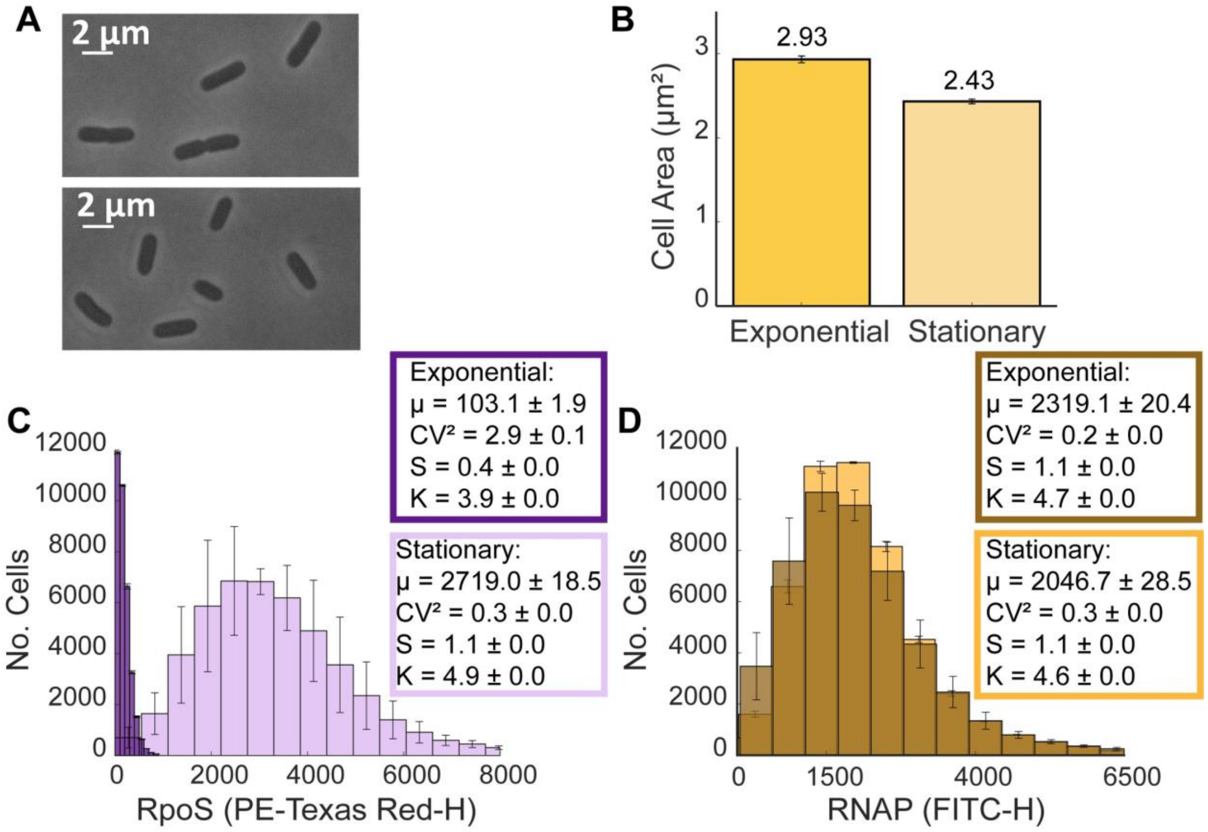


**Figure S3.** Morphology and physiology of cells and their respective RpoS levels when shifting from the exponential to the stationary growth phase. **(A)** Example phase contrast images of cells in the exponential (top) and stationary (bottom) growth phases. **(B)** Mean cell areas obtained from segmented phase contrast microscopy images (Section S2.3, main manuscript). Data from ~1500 cells per condition. **(C)** Single cell distributions of RpoS fluorescence intensity from MGmCherry cells obtained by flow cytometry in the exponential and the stationary growth phases. **(D)** Single cell distributions of RNAP levels in RL1314 cells measured by flow cytometry in the exponential and stationary growth phases. Also shown in **(C)** and **(D)** are the mean (µ), squared coefficient of variation ($CV^2$), skewness (S) and kurtosis (K) of the distributions, as well as error bars representing the standard error of the mean (SEM) from the 3 replicates.

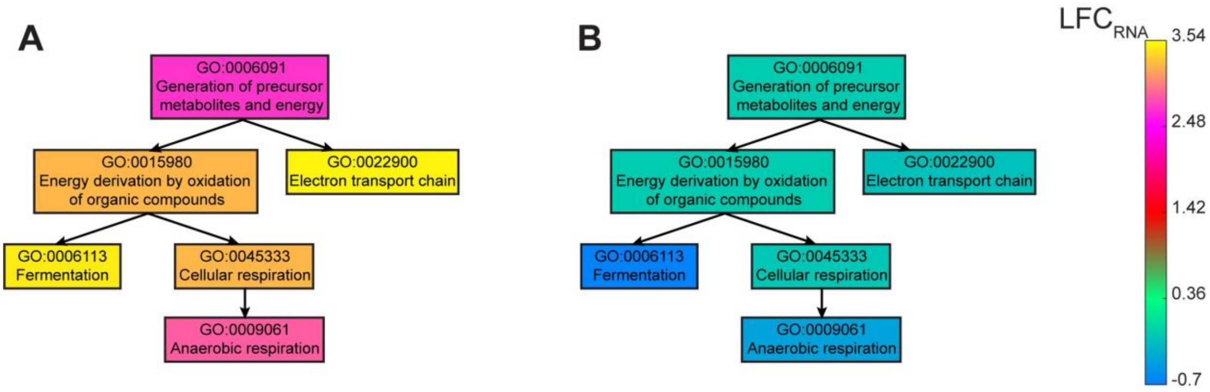


**Figure S4.** Overrepresented gene ontology terms for **(A)** $\sigma^{38+70}$ genes **(B)** non-$\sigma^{38+70}$ genes (Section 2.5, main manuscript) and the corresponding response strengths to the shift to stationary growth. Processes are organized from top to bottom as a function of the increasing specificity according to the gene ontology [14,15]. Each ontology is colored according to the mean LFC of the genes of the biological process.

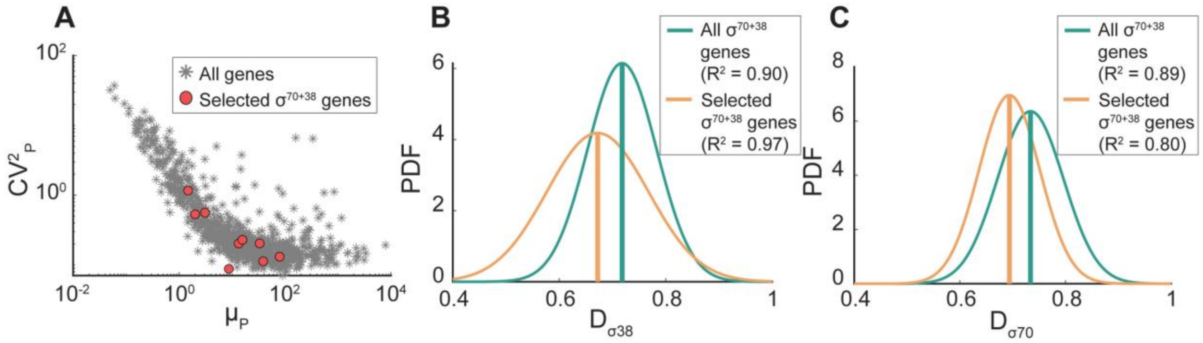


**Figure S5.** Genes selected to measure single-cell protein abundances. **(A)** Protein noise ($CV^2_P$) plotted against the mean protein numbers ($\mu_P$) according to [10]. The genes of interest are marked as red balls. **(B)** Gaussian fits to the distributions of $D_{\sigma 38}$ and **(C)** $D_{\sigma 70}$ of all $\sigma^{70+38}$ genes and of the selected $\sigma^{70+38}$ genes. Vertical lines mark the means of the distributions**.**

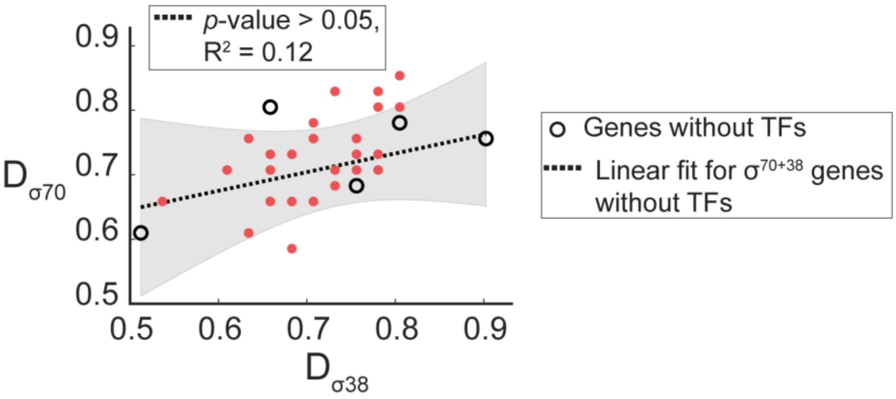


**Figure S6.** Scatter plot of $D_{\sigma 38}$ of a promoter sequence and $D_{\sigma 70}$ of the same promoter. The shadow of the best fitting line is the 95% confidence bounds. The $R^2$ value of the best fitting line and the p-value of the F test suggest that $D_{\sigma 70}$ and $D_{\sigma 38}$ are not significantly correlated.

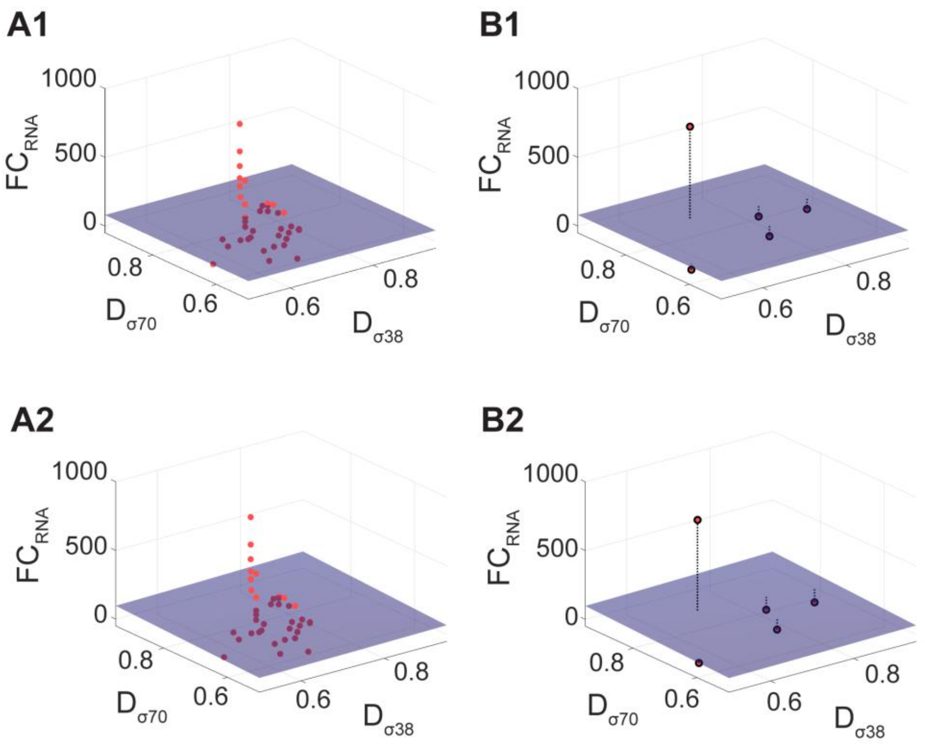


**Figure S7.** Fold changes in the RNA levels of $σ^{70+38}$ genes plotted against their promoter *p*-distances to the consensus sequence of $σ^{70}$ dependent promoters ($D_{σ70}$) and of $σ^{38}$ dependent promoters ($D_{σ38}$). **(A1-A2)** Best fitting surfaces to FC($μ_{RNA}$) assuming **(A1)** a linear function and **(A2)** a rational function. Only $σ^{70+38}$ genes with input TFs and FDR < 0.05 are included. Light red points are above the surface, while dark ones are below. **(B1-B2)** Same plots, with the surfaces obtained from (A1-A2) and applied to $σ^{70+38}$ genes without input TFs (FDR < 0.05). The dashed black lines depict the vertical distances between the estimated and measured FC($μ_{RNA}$).

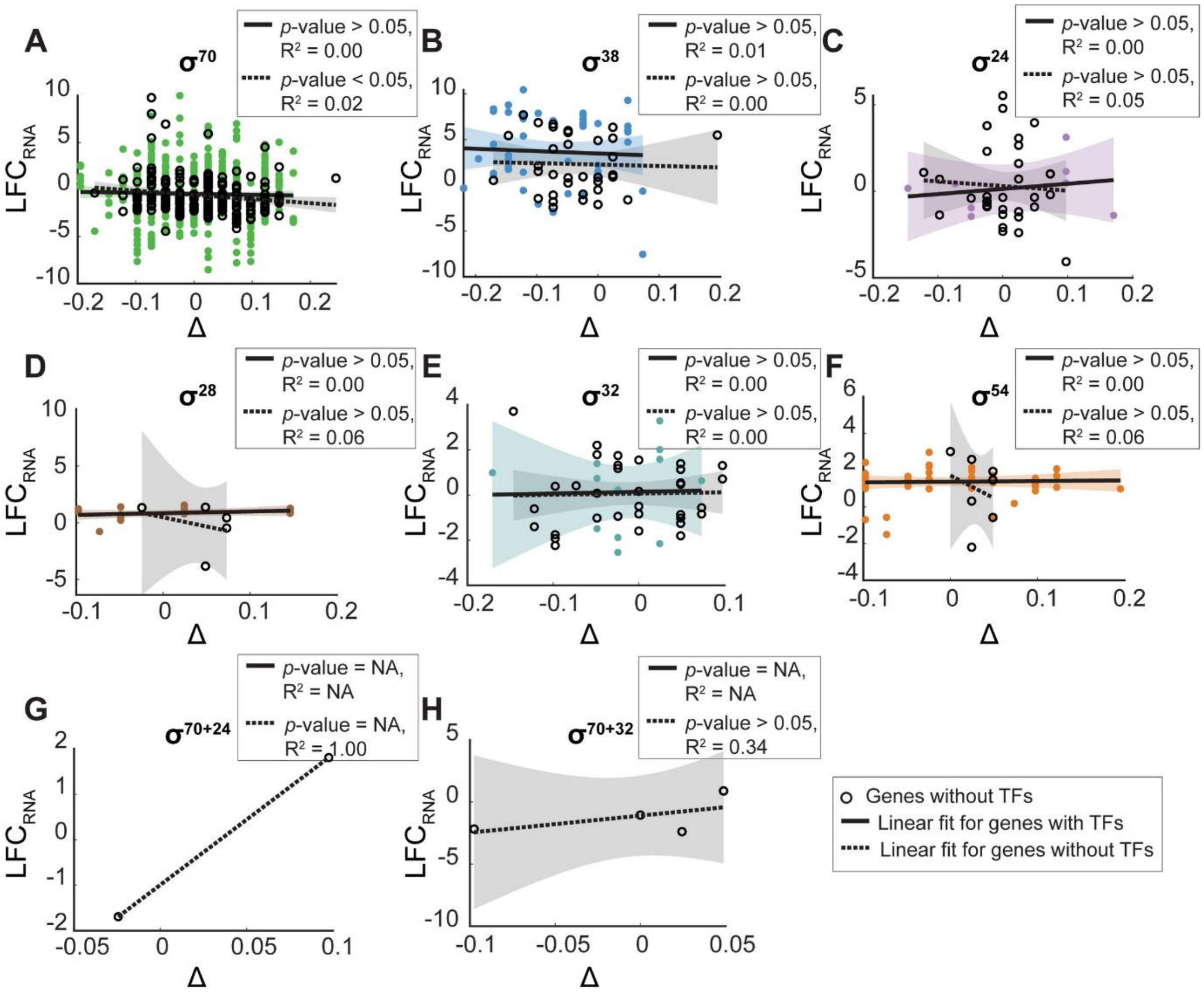


**Figure S8.** Fold changes in RNA levels and Δ. $LFC_{RNA}$ of genes with promoters with preference for **(A)** $\sigma^{70}$ **(B)** $\sigma^{38}$ **(C)** $\sigma^{24}$ **(D)** $\sigma^{28}$ **(E)** $\sigma^{32}$ **(F)** $\sigma^{54}$ **(G)** $\sigma^{70+24}$ and **(H)** $\sigma^{70+32}$ are plotted against $\Delta = D_{\sigma 38} - D_{\sigma 70}$. The shadows of the best fitting lines are the 95% confidence bounds.

### S3. Supplementary Tables

**Table S1.** Strains used to measure single-cell protein expression statistics [10].

| Strain name | Genotype | Source |
|---|---|---|
| SX1267 | F-, *Δ(argF-lac)169*, *gal-490*, *Δ(modF-ybhJ)803*, *λ[cI857 Δ(cro-bioA)]*, *alkA791-YFP(::cat)*, *IN(rrnD-rrnE)1*, *rph-1* | Yale CGSC (CGSC # 12822) |
| SX1506 | *F-, Δ(argF-lac)169, gal-490, Δ(modF-ybhJ)803, λ[cI857 Δ(cro-bioA)], dps-791-YFP(::cat), IN(rrnD-rrnE)1, rph-1* | Yale CGSC (CGSC # 13061) |
| SX1571 | F-, *Δ(argF-lac)169*, *gal-490*, *Δ(modF-ybhJ)803*, *λ[cI857 Δ(cro-bioA)]*, *IN(rrnD-rrnE)1*, *rph-1*, *pstS793-YFP(::cat)* | Yale CGSC (CGSC # 13126) |

| | | |
|---|---|---|
| SX1662 | *F-, Δ(argF-lac)169, gal-490, Δ(modF-ybhJ)803, λ[cI857 Δ(cro-bioA)], IN(rrnD-rrnE)1, rph-1, osmY792-YFP(::cat)* | Yale CGSC (CGSC # 13217) |
| SX1675 | F-, *Δ(argF-lac)169*, *gal-490*, *Δ(modF-ybhJ)803*, *λ[cI857 Δ(cro-bioA)]*, *IN(rrnD-rrnE)1*, *gor-791-YFP(::cat)*, *rph-1* | Yale CGSC (CGSC # 13230) |
| SX1767 | F-, *Δ(argF-lac)169*, *gal-490*, *Δ(modF-ybhJ)803*, *λ[cI857 Δ(cro-bioA)]*, *mlrA791-YFP(::cat)*, *IN(rrnD-rrnE)1*, *rph-1* | Yale CGSC (CGSC # 13322) |
| SX1791 | F-, *Δ(argF-lac)169*, *gal-490*, *Δ(modF-ybhJ)803*, *λ[cI857 Δ(cro-bioA)]*, *IN(rrnD-rrnE)1*, *rph-1*, *cpxR791-YFP(::cat)* | Yale CGSC (CGSC # 13346) |
| SX1793 | F-, *Δ(argF-lac)169*, *gal-490*, *Δ(modF-ybhJ)803*, *λ[cI857 Δ(cro-bioA)]*, *appC791-YFP(::cat)*, *IN(rrnD-rrnE)1*, *rph-1* | Yale CGSC (CGSC # 13348) |
| SX1960 | F-, *Δ(argF-lac)169*, *gal-490*, *Δ(modF-ybhJ)803*, *λ[cI857 Δ(cro-bioA)]*, *IN(rrnD-rrnE)1*, *rph-1*, *oxyR791-YFP(::cat)* | Yale CGSC (CGSC # 13515) |

**Table S2.** Plasmids from the Promoter fusion library [16] used to measure promoter activity by flow cytometry.

| Strain name | Genotype | Construction procedure | Source |
|---|---|---|---|
| MG1655+aidB | Same as MG1655; $P_{aidB}$-GFP, Kanr | Reporter plasmid $P_{aidB}$-GFP, Kanr in MG1655 | Promoter fusion library and this work |
| MG1655+asr | Same as MG1655; $P_{asr}$-GFP, Kanr | Reporter plasmid $P_{asr}$-GFP, Kanr in MG1655 | Promoter fusion library and this work |
| MG1655+pstS | Same as MG1655; $P_{pstS}$ -GFP, Kanr | Reporter plasmid $P_{pstS}$-GFP, Kanr in MG1655 | Promoter fusion library and this work |

**Table S3.** Number of genes transcribed by a single promoter with single and with dual σ preference.

| $\sigma^{70}$ | $\sigma^{54}$ | $\sigma^{38}$ | $\sigma^{32}$ | $\sigma^{28}$ | $\sigma^{24}$ | $\sigma^{19}$ | |
|---|---|---|---|---|---|---|---|
| 93 | | 64 | 9 | | 2 | | $\sigma^{70}$ |
| | 65 | 1 | | | | | $\sigma^{54}$ |

| | | | | | |
|---|---|---|---|---|---|
| 93 | | | | | **$\sigma^{38}$** |
| | 65 | | | | **$\sigma^{32}$** |
| | | 34 | | | **$\sigma^{28}$** |
| | | | 54 | | **$\sigma^{24}$** |
| | | | | | **$\sigma^{19}$** |

**Table S4.** Gene Ontology Overrepresentation Test. Tests done by PANTHER classification [12]) for **(A)** the cohort of $\sigma^{70+38}$ genes and **(B)** to the cohort of $\sigma^{38}$ genes. Listed are gene ontologies (GO) [14,15] related to biological processes overrepresented in the genes cohort (according to the Fisher's exact test with FDR correction). For each overrepresented ontology, we show the number of genes related to the specific ontology in the *E. coli* genome; the number of genes in the selected cohort related to the ontology; the number of genes expected to be present according to the size of the cohort; the fold-enrichment; the *p*-value of the Fisher's exact test to detect non-random associations between the two variables, and the false discovery rate (FDR). For FDR < 0.05, we reject the null hypothesis that there are no associations between the dual σ factor preference and the corresponding GO of the biological processes.

## A

| GO biological process | No. genes in genome | No. $\sigma^{70+38}$ genes | Expected No. genes | Fold Enrichment | *p*-value Fisher's exact test | FDR |
|---|---|---|---|---|---|---|
| Fermentation (GO:0006113) | 17 | 7 | 0.24 | 29.63 | $1.8\times10^{-8}$ | $5.8\times10^{-5}$ |
| Energy derivation by oxidation of organic compounds (GO:0015980) | 129 | 12 | 1.79 | 6.69 | $2.9\times10^{-7}$ | $2.3\times10^{-4}$ |
| Generation of precursor metabolites and energy (GO:0006091) | 210 | 14 | 2.92 | 4.80 | $1.2\times10^{-6}$ | $6.4\times10^{-4}$ |
| Anaerobic respiration (GO:0009061) | 61 | 10 | 0.85 | 11.80 | $2.6\times10^{-8}$ | $4.2\times10^{-5}$ |

| | | | | | | |
|---|---|---|---|---|---|---|
| Cellular respiration (GO:0045333) | 114 | 12 | 1.58 | 7.58 | $8.4\times10^{-8}$ | $8.9\times10^{-5}$ |
| Electron transport chain (GO:0022900) | 107 | 11 | 1.49 | 7.40 | $3.9\times10^{-7}$ | $2.5\times10^{-4}$ |

# B

| GO biological process | No. genes in genome | No. $\sigma^{38}$ genes | Expected No. genes | Fold Enrichment | *p*-value Fisher's exact test | FDR |
|---|---|---|---|---|---|---|
| Polyamine catabolic process (GO:0006598) | 11 | 5 | 0.23 | 22.17 | $1.07\times10^{-5}$ | $1.69\times10^{-2}$ |
| Putrescine metabolic process (GO:0009445) | 14 | 5 | 0.29 | 17.42 | $2.73\times10^{-5}$ | $2.86\times10^{-2}$ |
| Polyamine metabolic process (GO:0006595) | 18 | 6 | 0.37 | 16.26 | $5.59\times10^{-6}$ | $1.76\times10^{-2}$ |

**Table S5.** Coefficients of the Hill function fitted to the fold change in the mean protein levels over time relative to the level in the exponential phase ($FC_P$) of the genes pstS, aidB, and asr.

| Genes | Hill function $FC_P\left(t\right)=\frac{m\cdot t^s}{h^s+t^s}+b$ | | | |
|---|---|---|---|---|
| | Intercept *b* | Slope *s* | Half-activation *h* | Maximum *m* |
| pstS | 1.07 | 13.17 | 340.04 | 3.67 |
| aidB | 1.01 | 10.82 | 350.07 | 1.16 |
| asr | 1.00 | 5.58 | 338.97 | 0.52 |

**Table S6.** Goodness of fit, measured by the $R^2$ of the surface fitting of $FC_{RNA}$ as a function of $D_{\sigma38}$ and $D_{\sigma70}$. The surfaces were fitted to genes without input TFs and validated on genes with input TFs.

| Genes | Cohort | $R^2$ |
|---|---|---|
| $\sigma^{70}$ genes | Genes without input TFs | <0 |
| $\sigma^{38}$ genes | Genes without input TFs | <0 |
| $\sigma^{24}$ genes | Genes without input TFs | <0 |
| $\sigma^{28}$ genes | Genes without input TFs | <0 |
| $\sigma^{32}$ genes | Genes without input TFs | <0 |
| $\sigma^{54}$ genes | Genes without input TFs | <0 |
